# Polymorphic IGLV6-57 AL amyloid fibrils and features of a shared folding pathway

**DOI:** 10.1101/2024.12.04.626857

**Authors:** Parker T. Bassett, Binh A. Nguyen, Virender Singh, Patrick T. Garrett, James J. Moresco, Joshua Eddy, Shumaila Afrin, Maja Pękała, Christian Lopez, Bret M. Evers, Yasmin Ahmed, Justin L. Grodin, Lori R. Roth, Gurbakhash Kaur, Stephen Chung, Gareth J. Morgan, John R. Yates, Lorena Saelices

## Abstract

Immunoglobulin light chain (AL) amyloidosis is a systemic disorder caused by the misfolding and aggregation of free immunoglobulin light chains (LCs) secreted by abnormal plasma cells. The resulting amyloid fibrils deposit in multiple organs, leading to progressive dysfunction and increased morbidity and mortality. Despite recent advances, the molecular determinants of LC aggregation and phenotypic variability remain poorly understood. Structural characterization of *ex-vivo* fibrils provides key insights into these pathogenic processes. Here, we report cryo-electron microscopy structures of cardiac AL amyloid fibrils derived from an IGLV6 light chain. The fibrils display two distinct morphologies composed of single and double protofilaments, each adopting a previously unobserved fold. Comparison with previously reported AL fibril structures reveals that while individual mutations can alter the local conformation, IGLV6-derived fibrils share conserved structural motifs that may underlie common aggregation pathways. These findings expand the disease structural landscape and highlight sequence-dependent yet structurally constrained mechanisms of LC fibril formation.

## INTRODUCTION

Light chain amyloidosis (or AL amyloidosis) is a systemic amyloid disease that accounts for a significant proportion of patients diagnosed with systemic amyloidosis.^1–5^ Recent advances in detection methods and treatments have led to the identification of a greater number of cases and greater disease awareness. The disease is marked by the generation of an aberrant plasma clone, which freely secretes antibody light chain (LC) protein. Outside of the cell, these free LC proteins may misfold and aggregate into amyloid fibrils, eventually depositing in various organs and initiating the disease.^6,7^ AL can affect multiple organ systems, but commonly affects the heart and kidneys, and can be particularly aggressive, with some patients with advanced cardiac involvement only surviving for a few weeks after diagnosis.^2^ In conjunction, it is widely believed that AL amyloidosis is underdiagnosed due to poor recognition of red flag symptoms or symptom overlap with other diseases.^7^ Further innovations in diagnosis and treatment will alleviate this, necessitating study of the pathogenesis of the disease.

LC protein is a roughly 25 kDa protein secreted from plasma cells that most often exists coupled with a heavy chain protein within the larger antibody protein structure, ultimately consisting of two light chain and two heavy chain subunits per mature immunoglobulin G protein.^8^ Each LC subunit contains a variable light chain domain genetically fused to a constant light chain domain. The necessary diversity in antibodies arises from two processes, VJ recombination and somatic hypermutation.^6^ In LCs, VJ recombination shuffles together different combinations of variable and joining genes from the diverse set of germline-encoded genes.^6,9^ Then, somatic hypermutation selectively alters the LC sequence in three complementarity-determining regions (CDRs) which are ultimately responsible for epitope binding. The combination of these processes results in a diverse array of LC sequences and eventually a diverse array in amyloidogenic LCs between patients.

Although the amyloidogenic LC sequence varies from patient to patient, germline genes have been shown to correlate with organ involvement and clinical phenotype.^10^ A subset of genes is more frequently observed in AL amyloidosis clones tan in the healthy immune repertoire, and LCs derived from these genes are associated with amyloid deposition in different organs. The most common LC germline gene implicated in AL amyloidosis is IGLV6-57, which is predominantly associated with kidney, and to a lesser degree, heart involvement.^6,11–13^ These phenotypic trends are likely driven by the unique features present in a gene sequence. Studies seeking to explore this role have focused on the structural conversion from a natively folded LC, to a partially or fully unfolded monomer, and finally formation of the misfolded amyloid fibril. Recently, cryo-EM studies have shown a diverse range of *ex-vivo* AL fibril structures derived from various germline genes.^14–21^ These fibril structures demonstrate a complete loss of the native structure and adoption of diverse amyloid folds. Only a few objective similarities exist, such as inclusion of the variable domain in the fibril core and an intact, though reversed, disulfide bond. However in spite the structural diversity, it has been hypothesized that AL fibrils derived from the IGLV3-1 and IGLV3-19 genes, each with high degrees of LC sequence similarity, may be structurally related.^18,20^ Although not yet confirmed, this observation suggests that LC sequences derived from genes similar in sequence may lead to the formation of structurally related fibrils despite somatic variability.

Here we report new cryo-EM structures of AL fibrils, extracted from the explanted heart of a patient with AL amyloidosis, derived from the IGLV6-57 gene. These fibrils display multiple morphologies with single or double protofilaments, one of which presents with an uncommon rotational symmetry, raising new questions about the fibril formation process in AL amyloidosis. The comparison of these new structures to existing IGLV6-57 fibril models contributes to the understanding of the mechanism of aggregation of AL fibrils with similar sequences.

## RESULTS

### Extracted fibrils from an explanted heart are made up of a truncated **λ** light chain protein

We sourced cardiac tissue from the explanted heart of a patient at UT Southwestern Medical Center diagnosed with AL amyloidosis characterized by heart and kidney involvement. Upon diagnosis, histopathological analysis demonstrated positive Congo red staining in the right ventricular endomyocardium. Fibrils were directly visualized by electron microscopy. Further clinical testing revealed the patient had AL amyloidosis due to a λ-class LC, the more common of the two main λ or κ LC isotypes.^6^ The patient was treated with cyclophosphamide, bortezomib, dexamethasone, and daratumumab followed by an autologous stem cell transplantation, resulting in complete hematological response. Over a year and a half after stem cell transplantation, the patient underwent dual-organ transplantation for advanced heart and kidney failure. Following informed consent, we received the explanted heart immediately following the surgery and sectioned the tissue for further histology and fibril extraction. The histological studies of fixed tissue sections revealed amyloid deposits throughout the heart, including the left and right ventricles (Fig. 1a). Amyloid fibrils were extracted from unfixed tissue using a water-based extraction procedure.^22,23^ Transmission electron microscopy (TEM) confirmed the presence of fibrils after extraction (Fig. 1b and Supplementary Fig. 1a).

**Figure 1.**
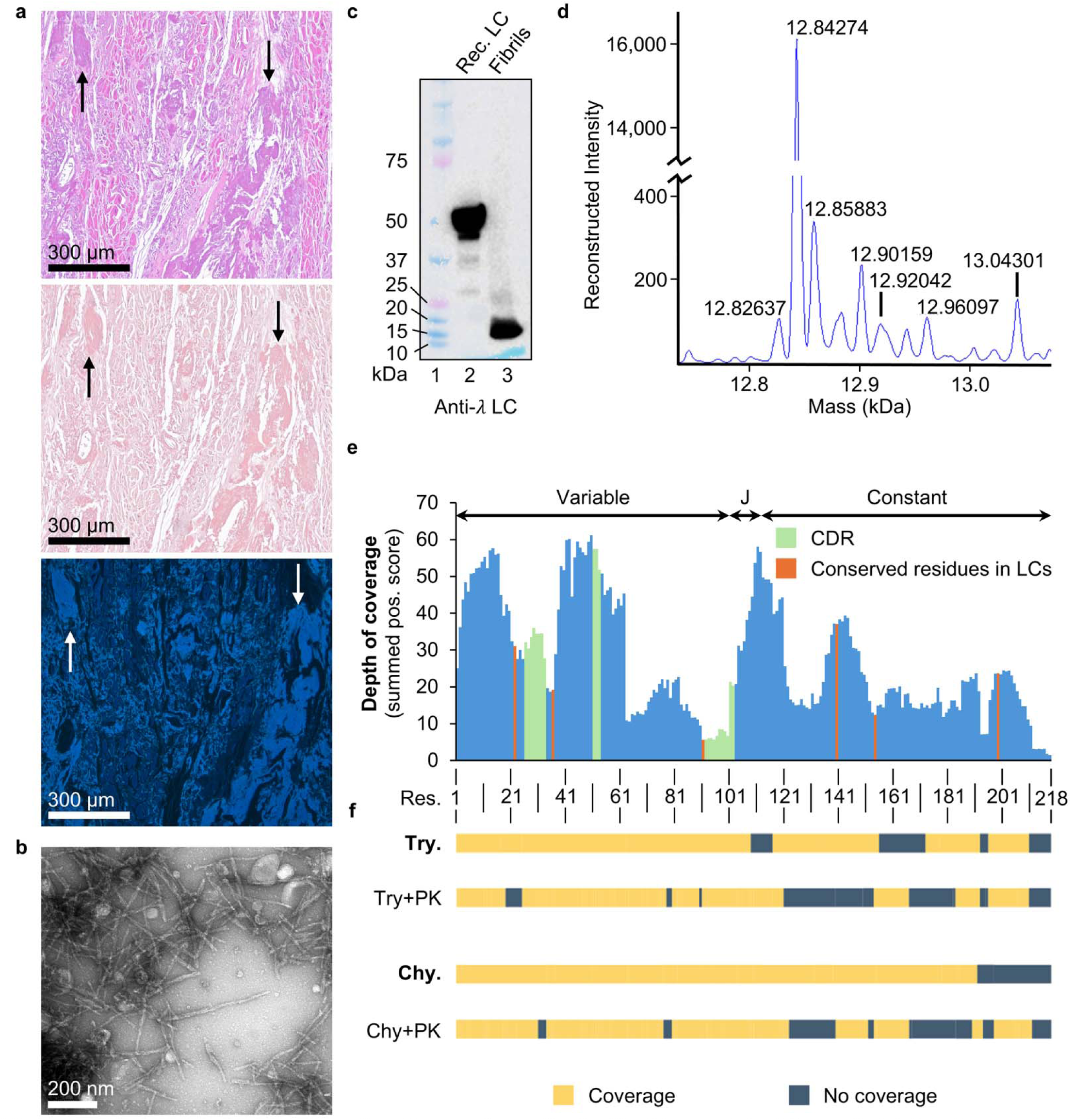
Fibrils extracted from cardiac tissue of a patient with AL consist of truncated λ LC fragments. **a)** Histology slides from the patient’s right ventricle of the heart, stained with hematoxylin and eosin (top), Congo red (middle), and thioflavin S (bottom). Amyloid deposits marked by arrows. 1Ox nominal magnification, scale bar 300 µm. **b) TEM** of fibrils extracted from cardiac tissue. 20,000x nominal magnification, scale bar 200 nm. **c)** Western blot under reducing and denaturing conditions with anti-λ LC antibody targeting a recombinant LC protein control (lane 2) and fibrils extracted from the patient sample (lane 3). The positive control was purchased commercially (Abeam) and runs as a dimer ~50 kDa on SOS-PAGE. **d)** Intact mass analysis shows the dominant protein species in the extracted fibril solution to have a mass of 12.8 kDa, approximately similar to the band in lane 3 of panel 1c. **e)** Short peptide reads from enzymatic digestion were mapped to the predicted sequence of the AL fibrils and matching residues were scored. The resulting score demonstrates the depth of coverage on our predicted sequence. CDRs are shaded in green and conserved residues (such as cysteines) are shaded in orange. **f)** Additional digestion of fibrils using proteinase K (PK) revealed regions in our LC sequence that are likely found outside the structured core of the AL fibrils, represented by areas with no sequence coverage after treatment with PK. Try. - Trypsin, Chy. - Chymotrypsin.

To further validate the identity of the amyloid extract, we performed western blot and mass spectrometry (MS). In the western blot, we observed the presence of a predominant λ LC species between 10 and 15 kDa, indicating the presence of truncated light chains (Fig. 1c).Intact MS analysis showed a predominant protein species with a mass of 12.8 kDa, consistent with the size of the λ-light chain band observed in the western blot data (Fig. 1d).

### Mass spectrometry indicates fibrils are made up of IGLV6-57 derived light chains

Sequence information was unavailable for this patient, so neither the clonal protein sequence involved in the amyloid, nor the germline genetic identity was known prior to structural determination. In order to determine the amino acid sequence of the fibrils, we used an MS-based approach with the software Casanovo and Stitch.^24–27^ Denaturation and digestion of the purified *ex-vivo* fibrils with trypsin, chymotrypsin, and elastase yielded spectra that were used in a *de-novo* sequence determination (Fig. 1e). Thus, we determined that the fibril sequence is derived from either the IGLV6-57*03 or IGLV6-57*04 alleles, later supported by the cryo-electron microscopy (cryo-EM) data as described below. The constant region was predicted to be IGLC2, and the joining region was predicted to be IGLJ2. However, the latter assignment is ambiguous since the sequences of joining regions IGLJ2 and IGLJ3 are indistinguishable at the amino acid level. Through this process, we were able to determine amino acid substitutions and insertions not encoded in the germline precursor genes, which are discussed in more detail below.

As an additional validation of this *de novo* sequence, we analyzed the fibril extract by MS before and after proteinase K digestion. The MS data analysis prior to proteinase K digestion identifies many proteins present in our sample, including amyloid-signature proteins such as apolipoprotein E, apolipoprotein A, serum amyloid P, and several forms of collagen, including collagen VI (Supplementary Data 1). Since fibril cores have been shown to be proteolytically resistant even to proteinase K, we hoped that this technique would enrich LC sequence regions involved in the fibril core within the MS data.^28^ Expectedly, a search of a database including our *de novo* sequence revealed reduced spectral counts in sequence regions predicted outside the fibril core, and vice versa (Fig. 1f).

### A high-resolution density map reveals the amyloid fold of IGLV6-derived light chains

Using our validated extracted fibrils, we prepared cryo grids using a Vitrobot Mark IV plunge freezer and screened them using a Talos Arctica 200 kV electron microscope. Several rounds of optimization were needed to determine the grid freezing conditions that yielded a suitable concentration of evenly distributed fibrils frozen in vitreous ice. Grids with optimal fibril concentration, good quality ice, and dispersed fibrils were saved for data collection, to be performed at Stanford-SLAC Cryo-EM Center.

Cryo-EM screening confirmed the presence of diverse configurations of fibrils, suggesting multiple morphologies (Fig. 2a and Supplementary Fig. 1b). Particle picking using Topaz resulted in over 1.4 million particles, and their 2D classification using RELION revealed single and double protofilament morphologies (Supplementary Fig. 2a). We selected 2D classes for both single and double morphologies and stitched them manually to determine their approximate fibril crossover length (Fig. 2b, top, and Supplementary Table 1). We were unable to generate a suitable initial model in RELION for the single filament morphology and so we used a featureless cylinder for subsequent 3D classification. In contrast, we were able to generate an initial model for the double protofilament morphology which was used for further processing (Fig. 2b, bottom, Supplementary Fig. 2b). During the 3D classification, we only observed the two unique structural classes corresponding with one and two protofilaments. (Fig. 2c). The Supplementary Fig. 3 schemes the overall helical reconstruction process.

**Figure 2.**
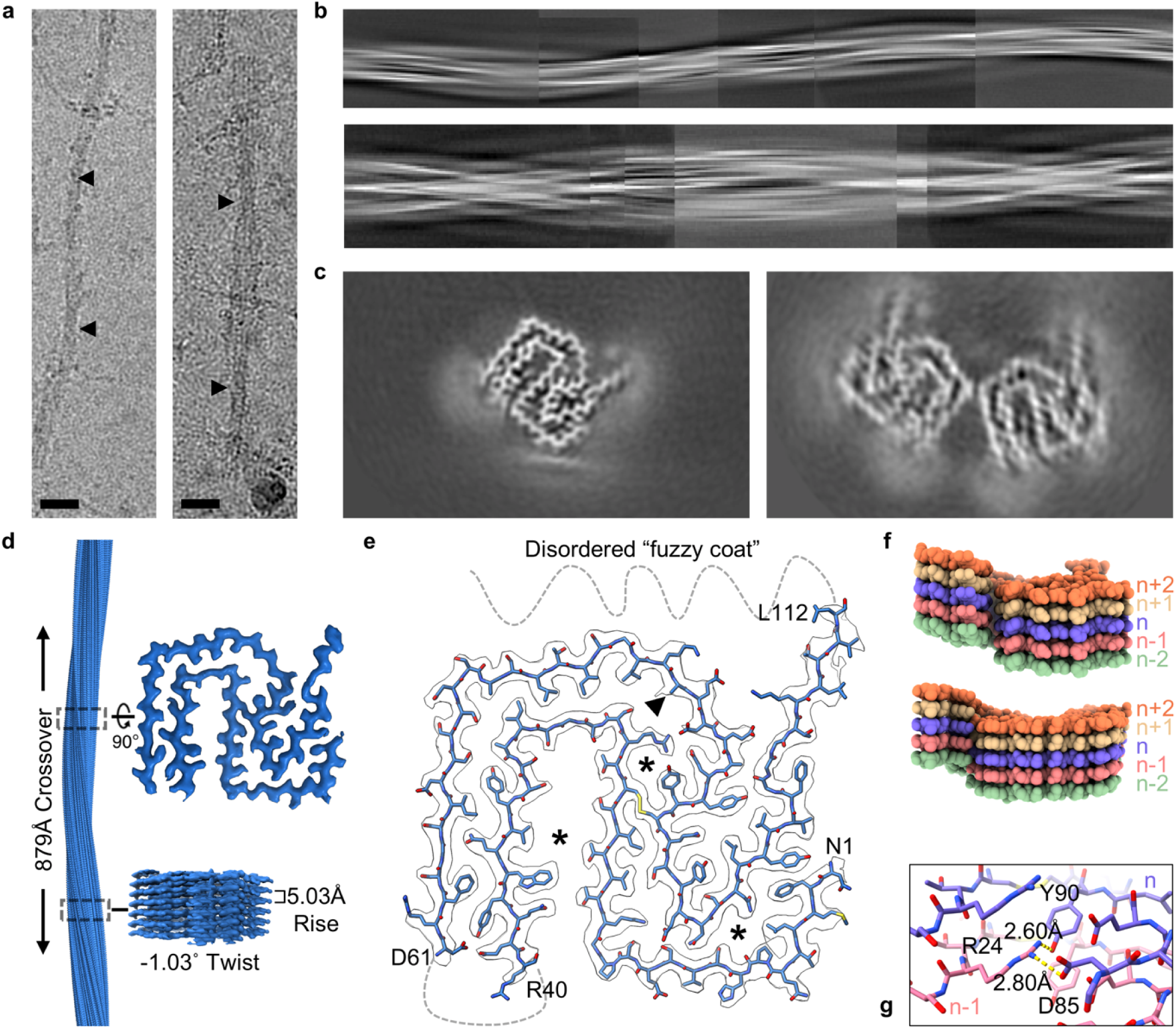
Cryo-EM of cardiac AL fibrils reveals polymorphism and interactions between fibril layers. **a)** Cryo-EM images show both single (left) and double (right) protofilament structures. Crossover lengths are marked by triangles. 105,000x nominal magnification, scale bar 40 nm. **b)** Full-length crossovers were assembled from representative 2D classes. 2D classes showed both single (top) and double (bottom) protofilament morphologies. **c)** 30 classification revealed two dominant morphologies: a single protofilament structure (left), and a double protofilament structure (right). d) Fibril topology, including helical rise, twist, and crossover. The density map was solved to a resolution of 3.0Å. **e)** The fibril model consists of a double layered protein spanning residues 1-40 and 61-112. Asterisks (*) mark unmodelled densities. Triangle symbol (▲) marks residue 83, with unknown identity, modelled as an alanine in the structure. f) A single layer of the fibril is spatially distributed along the fibril axis and not planar. The top view shows parts of the fibril that interact one layer apart. The bottom view shows parts of the fibril that interact across two layers. **g)** An example of interactions between layers of the fibril. Shown are Tyr 90 and Asp 85 which closely interact with Arg 24 in the layer below at distances approximately 2.60 and 2.80Å.

The single protofilament morphology was the most abundant in the 2D and 3D class averages. We solved the structure at a resolution of 3.0 Å based on the gold-standard FSC curve (Supplementary Fig. 3c). At this resolution, the cross-beta fibril structure was clearly resolved, and the overall conformation of the fibril core was determined. Local resolution of the map is best at the center of the core at 2.8Å, and some areas of the core were less resolved and more likely to be disordered (Supplementary Fig. 3b). Additionally, we observed three disconnected densities unlikely to result from sidechain orientation. (Fig. 2e, asterisks, Supplementary Fig. 4a).

We then used the MS-predicted sequence to model the atomic structure (Fig. 2e). The fibril core consisted of two LC regions, comprising residues Asn 1 to Arg 40 and Asp 61 to approximately Leu 112 (based on consecutive residue numbering), which would encompass CDRs 1 and 3, respectively. The resolution of the structural data is not high enough to completely resolve the conformation of the residues 108 to 112, so the side chain conformations are modelled with less confidence. Based on the previous WB and MS data, we hypothesize that the truncated LC would extend to residue 120, though we did not observe density beyond residue 112, suggesting that any amino acids in this region are disordered and flexible. Additionally, the region between residues 41 and 60, encompassing CDR 2, appeared disordered and/or absent. All these disordered residues would comprise the so-called “fuzzy coat” of the AL fibrils. Consistently, our MS analysis after proteinase K digestion showed that the C-terminal residues are sensitive to degradation (Fig. 1f). Additionally, gel electrophoresis under reducing conditions after heat denaturation of the proteinase K digested fibrils revealed the appearance of two species with a molecular weight of ∼12 kDa and ∼10 kDa (Supplementary Fig. 5). These masses correspond to the mass of the fibril core up to near residue 120, with or without the disordered residues 41 to 60 of the fuzzy coat.

### Modeling reveals interactions that overcome a low predicted fibril stability

We computationally estimated the solvation energy of the IGLV6 fibrils described here based on the buried surface area using previously published methods.^29^ Although the estimated solvation energy of fibrils appears to be the lowest of all *ex-vivo* AL fibril structures available in the Amyloid Atlas to date (Supplementary Fig. 6b,c), we found that the single protofilament morphology is stabilized by multiple interactions. First, the structure of the fibril core in the single protofilament morphology is centered around the intact disulfide bridge, linking Cys 22 to Cys 91. Second, each monomer of the fibril adopts a structure that is not planar, instead interlocking with the monomers above and below along the fibril axis. This creates opportunities for side chains to form interactions with immediately adjacent monomers (Fig. 2f). One example of this is shown in Figure 2g, where Arg 24 of one monomer interacts with Asp 85 and Tyr 90 in a separate monomer. Moreover, residues near the C-terminal end of the fibril core are close enough to interact with residues from a monomer two layers away (Fig. 2f). The fibril structure displays further electrostatic, hydrophobic, and polar interactions (Supplementary Fig. 4b). Finally, we observe the presence of unmodelled densities, possibly waters, regularly spaced alongside the layers of the fibril and in close proximity to polar residues (Supplementary Fig. 4a).

### Double protofilament fibrils have a unique symmetry with a small interaction surface

We further analyzed the additional double protofilament morphology observed during 3D classification using RELION (Fig. 2c). This double protofilament morphology has a rare rotational symmetry, with two similar protofilaments related by a 180° rotation perpendicular to the fibril axis (Fig. 3a). When compared to the single protofilament morphology, the double protofilament morphology had a distinct helical rise and twist value (Supplementary Table 1). This resulted in a different helical crossover length compared to the single protofilament morphology (Fig. 3b). Based on FSC curves, the double protofilament morphology density map had a resolution of 3.78Å (Supplementary Fig. 3c). In spite of the reported resolution, the map quality was insufficient to directly model the structure. However, since the 3D class shares a structural resemblance with the single protofilament morphology, we used molecular docking to fit the single protofilament model into the double protofilament map (Fig. 3b). The docking resulted in a visually reasonable fit and indicated that Asn 72 and Ser 73 are the residues involved in the protofilament interface.

**Figure 3.**
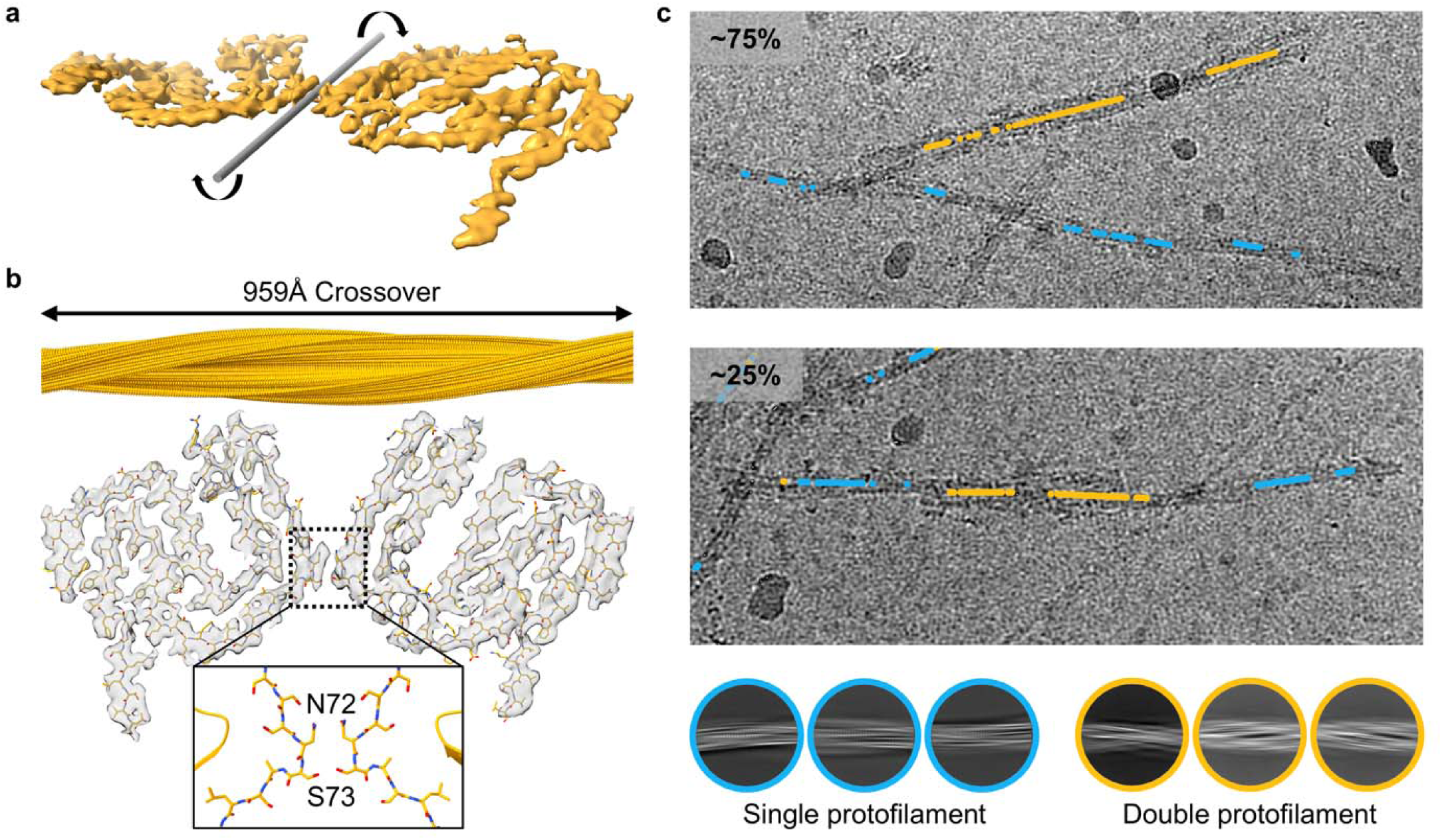
AL fibrils can adopt structures with multiple protofilaments that exist as isolated fibrils or within the same fibril. **a)** The double protofilament structure adopts a rare mode of symmetry, where the grey rod represents the rotational axis. **b)** The double protofilament structure is presented with a rigid-body fit of the single protofilament structural model. The inset shows the residues nearest the possible interaction site. **c)** Tracking the location of different polymorphs in the raw micrographs using RELION particle coordinates revealed where each polymorph appears in relation to the other. Blue dots represent single protofilament morphologies and yellow dots represent double protofilament morphologies. The majority of micrographs show the polymorphs isolated from each other (top), but a small minority of micrographs reveal the two polymorphs co-existing on the same fibril (bottom).

### Particle tracing reveals the coexistence of polymorphs on the same fibril

We then set out to determine how the single and double protofilament morphologies are distributed across the cryo-EM grid and how they might be interacting. To do this, we grouped particles from the 2D class averages that showed both single and double protofilament morphologies. Each group contained particles with an estimated resolution equal to or better than 5Å, resulting in 388,527 and 341,845 particles, respectively. This cutoff ensured that these particles were sourced from the two structures reported here, and did not belong to other undetermined morphologies, should there be any. We then leveraged RELION’s particle coordinate data to directly correlate the position of the particle in the raw micrograph to the protofilament morphology that it belongs to. The majority of fibrils were represented by one polymorph, and thus composed entirely of only one or two protofilaments (Fig. 3c). However, a minor population of fibrils were observed with both polymorphs co-existing on the same total fibril (Fig. 3c). Additional representative tracing images are found in Supplemental Fig. 7.

### Specific sequence regions and amino acid substitutions are likely to drive aggregation

To understand the potential mechanism of aggregation leading to these IGLV6 fibrils, we computationally predicted aggregation-prone regions (APRs) in their LC sequence. We used ten APR prediction methods via the AmylPred2 server, and calculated an overall APR prediction for each residue with a score from zero to ten, reflective of how many methods positively predicted each residue to be part of an APR.^30–36^ Our analysis showed many areas predicted to be APRs across the entire LC sequence (Fig. 4). Within the fibril core, strongly predicted APRs are centered around a few key areas. First, each Cys residue is involved in an APR, which has previously been experimentally verified.^37^ Second, strongly predicted APRs containing hydrophobic or aromatic residues are found at residues 33-38, 74-79, and 98-104. The APRs at residues 33-38 and 98-104 overlap with CDRs 1 and 3, which have also been previously identified to be hotspots for experimentally determined APRs in IGLV6-derived LCs.^37^ One of the APRs outside the fibril core was observed at residues 46-51, which overlaps CDR 2. CDR 2 is absent from the fibril core of the 9ELS structure as well as the two previously published 6HUD and 9OKA structures.^14,21^ However, CDR 2 is completely or partially observed in the fibril core of all IGLV3-derived^17,18,20^ and IGLV1-derived^15,16^ AL fibril structures to date. What role APRs near CDR 2 play in the aggregation of AL fibrils is yet to be determined. The other strongly predicted APR was observed at residues 137-154. This APR has previously been studied in the context of differential proteolysis between dimeric free LCs and natively folded immunoglobulin G proteins^38^. More experimental work is needed to understand the true role all of the APRs in IGLV6-derived LCs play in AL fibril formation.

**Figure 4.**
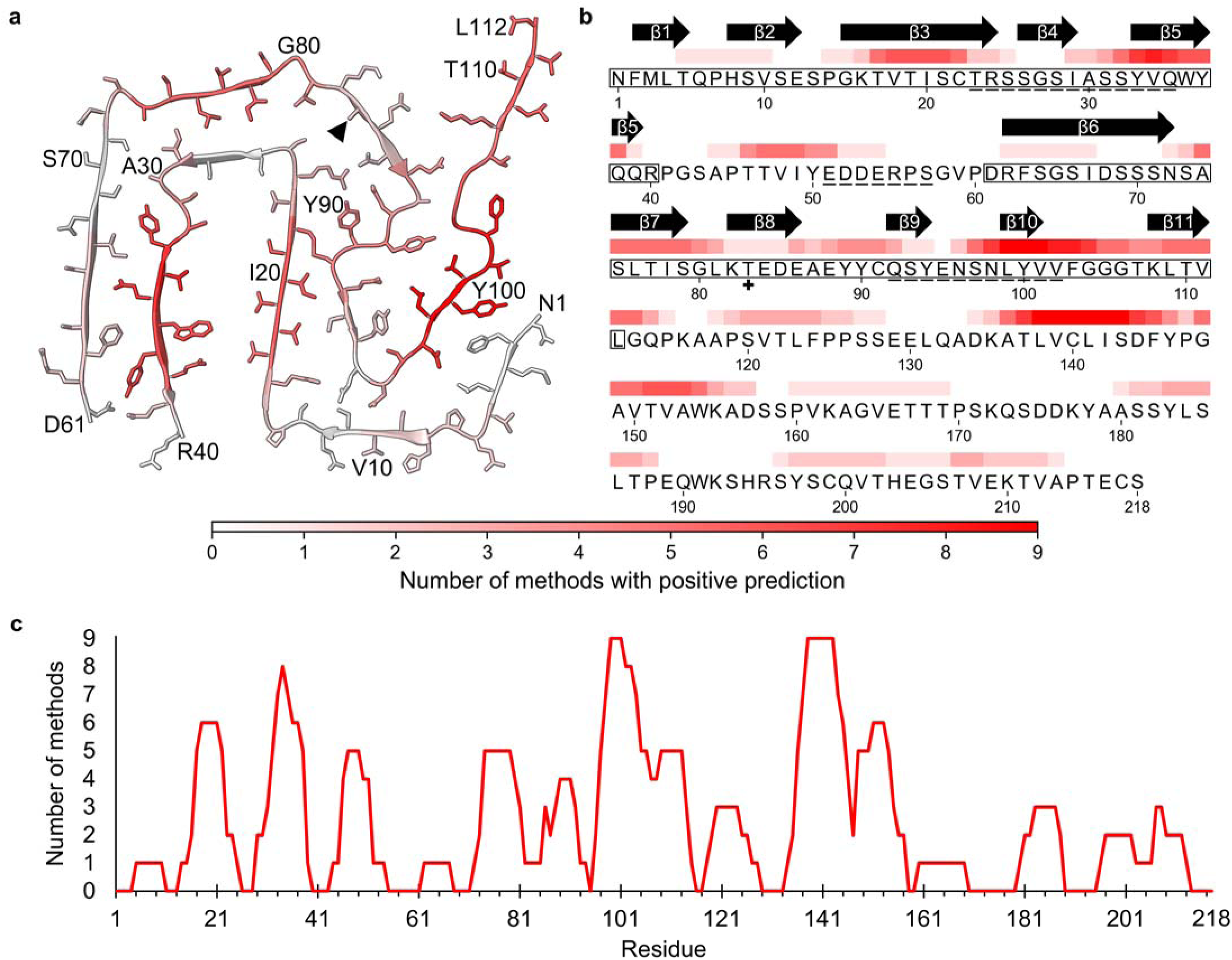
Predicted APRs reside in the structured core of the fibril. **a)** A heat map showing the predicted APRs in the AL fibril structure. **b)** Sequence identity of the predicted APRs expressed as a heat map, with secondary structure shown above the corresponding residues. Residues in the core of the fibril are boxed, and CDRs are underlined. For panels **a)** and **b),** darker red indicates a higher number of methods included in AmyIPred2 predict a residue as part of an APR. Triangle symbol (▲) marks residue 83, with unknown identity, modelled as an alanine in the structure and reverted to the germline residue assignment in the sequence. c) A line graph showing the APRs and number of methods used by AmyIPred2 that predict a residue to be part of an APR.

We sought to analyze how the residues that differ from the germline sequence, some of which reside in the core of the fibril, may contribute to the fibril stability (Fig. 5). We found that none of the mutations located in the structure core contribute strongly to the estimated solvation energy of the fibril (Supplementary Fig. 6b). It is also worth noting the presence of arginine at position 24 (position 25 according to Kabat numbering). The presence of an arginine at this position may affect the native protein stability and/or fibril formation rate, as suggested in previous studies.^11^ Though we observe no effects of these differences based on our energetic analysis, specific residues likely play a more subtle role in stabilizing the fibril structure, as we discuss later.

**Figure 5.**
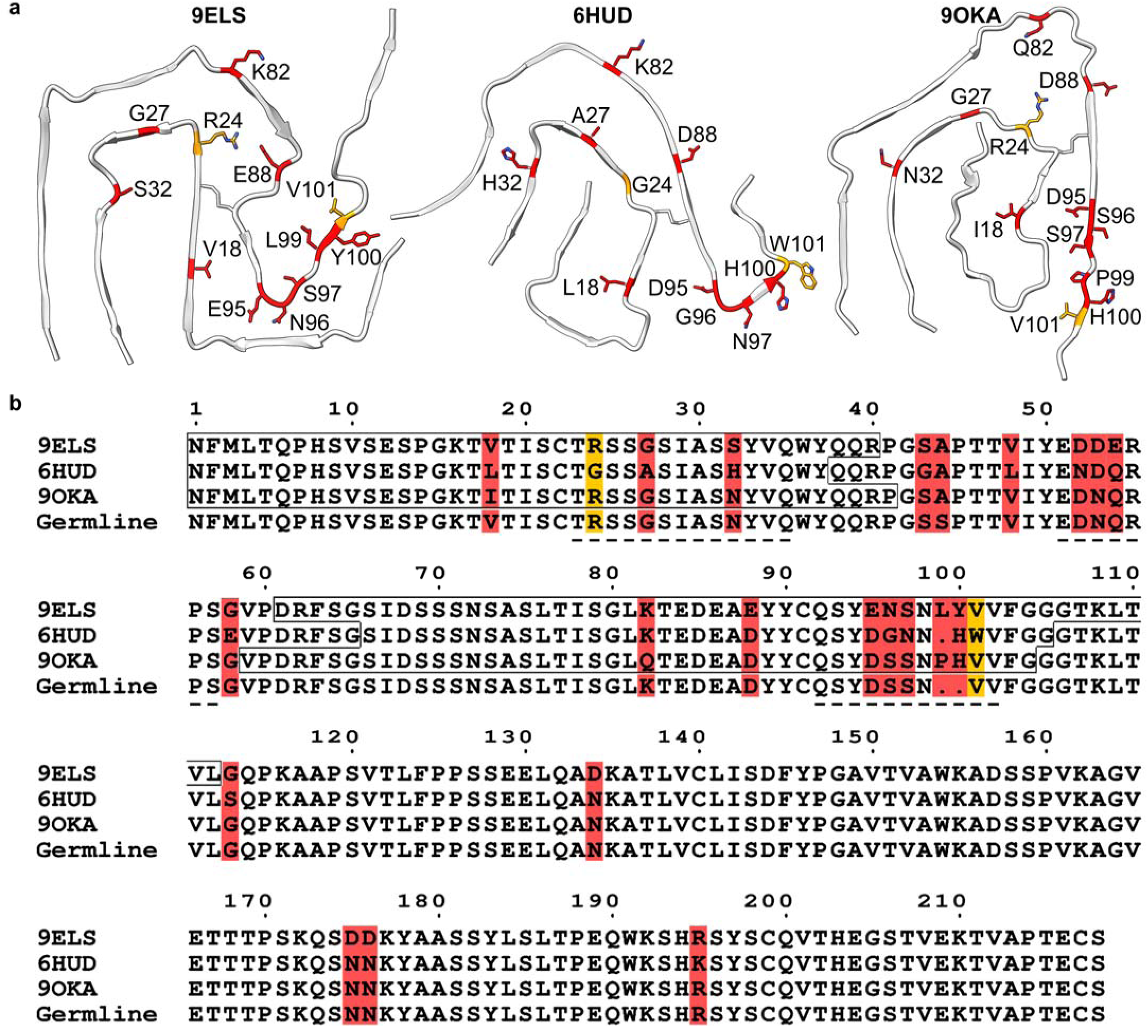
The sequences of previous AL fibril structures are highly similar to the 9ELS AL fibrils. The 9ELS, 6HUD, and 9OKA AL fibril structures **(a)** and sequences **(b)** are shown with their amino acid differences from both each other and the combined IGLV6-57, IGLJ2, and IGLC2 germline genes. Residues that differ between the four aligned sequences are highlighted in red. The 6HUD structure has a sequence derived from a different IGLV6-57 allele and thus varies at residue 24 and 101, which are highlighted in orange. Residues in the core of the fibril are boxed, and CDRs are underlined. The sequence numbering is sequential and based on the 9ELS sequence.

With two other IGLV6-derived AL fibril structures available, we compared the sequence between our structure and the others, which we refer to as their PDB codes 9ELS (the structure reported here), 6HUD^14^, and 9OKA^21^. We used the Needleman-Wunsch pairwise alignment algorithm to first determine the similarity and exact identity between the fibril sequences for 9ELS, 6HUD, and 9OKA.^39–41^ We compared both the full length sequences of each amyloidogenic LC, as well as only those residues involved in each fibril core and including the disordered loops encompassing CDR2. The first 112 residues of our structure 9ELS had an 80.4% sequence identity and an 87.5% sequence similarity when compared to 6HUD, and an 83.9% sequence identity and a 92.0% sequence similarity when compared to 9OKA. Specifically, within the core of the fibril, these differences amount to 11 different amino acids between 6HUD and our structure, and 8 different amino acids between 9OKA and our structure (Fig. 5). The sequence identity and similarity values were similar or higher when comparing full-length LC sequences.

## DISCUSSION

Cryo-electron microcopy has helped uncover the structures of AL fibrils showing an unprecedented sequence-specific structural diversity.^14–20^ In this study, we extracted *ex-vivo* AL amyloid fibrils from a patient with AL amyloidosis, which were made up of LCs derived from the IGLV6-57 germline gene (Fig. 1). We determined the structure of the fibrils and observed two unique fibril morphologies (Fig. 2). Ours is the first report showing the cryo-EM structure of AL fibril polymorphs with different numbers of protofilaments. We determined the structure of a dominant single protofilament morphology at 3.0 Å resolution and described a double protofilament morphology (Fig. 2 and 3). Both structures share the same protofilament fold with differences in the number of protofilaments found in the fibril. *De novo* sequencing by MS enabled us to generate a structure model for the dominant morphology (Fig. 1 and Fig. 2). We also used docking and fibril tracing to gather additional structural insights, as discussed below (Fig. 3).

Our present structure displays a few similarities to past AL fibril structures.^14–21^ All AL fibril structures have an intact disulfide bond that links two beta strands oriented antiparallel to each other. This contrasts with the native structure, where the beta strands linked by the disulfide bond are oriented parallel to each other. Additionally, the structured core of the fibrils described here includes only the variable and joining region, which is consistent with most of the published AL fibril structures to date^14–19,21^ and other earlier analyses.^42–44^ These AL fibrils also lack ordered density for CDR 2, which is consistent with some other AL fibril structures.^14,16–19,21^ When comparing the predicted solvation energy of each layer of the AL fibrils, our structure and other IGLV6-57-derived fibrils require the least amount of energy to solvate of the AL fibril structures to date (Supplementary Fig. 6c). Lastly, our structure also consists of fibril monomers with interlayer nonplanar interactions through side chains and backbone, similar to previous AL fibril structures.^14,21^

The most striking difference between our data and nearly all previously determined cryo-EM AL fibril structures is the presence of polymorphic species. To date, multiple morphologies have been observed in cardiac amyloid fibrils from other amyloidoses.^45,46^ However, all but one of the many published cryo-EM studies show AL fibrils with a single protofilament morphology.^14–20^ The one exception shows single and double protofilament morphologies, and describes the structure of the dominant single morphology.^21^ We found the presence of IGLV6-57 amyloid fibrils with a double protofilament morphology (Fig. 2), and structural docking allowed us to visualize the molecular nature of these fibrils and approximate the surface where these protofilaments associate (Fig. 3b). While the surface between protofilaments is small within a single layer of a fibril, the small surface is compounded across the length of an entire fibril, resulting in a favorable interaction surface.

This double protofilament morphology represents the first cardiac amyloid fibril with 2-fold rotational symmetry perpendicular to the fibril axis (Fig. 3a). In fact, across the 615 publicly available amyloid fibril structures in the Amyloid Atlas,^29^ there are only 9 fibril structures with a similar rotational symmetry perpendicular to the fibril axis, of which only 4 are *ex vivo* fibrils, all derived from brain tissue.^47–53^ This unique rotational symmetry of protofilaments raises questions about how they undergo elongation and assembly. Previous studies on insulin and Aβ42 amyloid fibrils have shown fibrils tend to exhibit polarity with different elongation rates at their “plus” and “minus” ends.^54,55^ Given that each of the protofilaments in the double protofilament morphology reported here are oriented in an antipolar fashion, each of their tips expose both “plus” and “minus” sites, and therefore, both “fast” and “slow” elongation sites (Supplementary Fig. 8). This is supported by the particle tracing data, in which we observe the presence of single protofilaments extended from double protofilament morphologies (Fig. 3c). As for the assembly of the double protofilament morphology, we speculate that there are at least three possibilities. First, the formation of a second protofilament occurs by the secondary nucleation of LCs onto the surface of a pre-existing single protofilament morphology that then elongates along the available fibril lateral surface. Second, single protofilaments nucleate and elongate in isolation before wrapping around each other thereby associating into the double protofilament morphology. Lastly, both sites within the same fibril end of a double protofilament elongates at the same time but at different rates. Our data are not sufficient to support or eliminate these models. To completely define the mechanisms by which these polymorphs form and mature, further study is warranted.

A closer comparison of our structure 9ELS with two previously determined *ex-vivo* IGLV6-57 structures allows us to contrast AL fibrils of the same genetic precursor. Previously, it has been postulated that AL fibrils from a common genetic precursor subfamily have a related fold.^20^ This was based on observations of similar fibril folds from structures of AL fibrils originating in separate organs of a single patient as well as separate patients with highly similar AL fibril sequences from the same genetic precursor subfamily.^18–20^ Our structure and the other IGLV6-57 structures reveal a similar trend. At a sequence level, the amyloidogenic LCs we describe here are highly similar to the previously published 6HUD and 9OKA fibril structures, though still display some structural differences (Fig. 5).^14,21^

The specific differences between the three structures could be attributed to differences in amino acids and the interactions and conformations adopted by them. Between our structure 9ELS and the 6HUD and 9OKA structures, there are 11 and 8 residue substitutions in the core of the fibrils, respectively (Fig. 5). Interestingly, compared to the germline, all three sequences have untemplated insertions between the variable and joining gene segments, with one insertion in 6HUD and two insertions in 9ELS and 9OKA. In our structure, but not in the other two, the insertion of Tyr 100 allows for possible π-π interactions with Phe 2 and perhaps explains the differential folding of the N-terminus on the periphery of the fibril core.

A specific residue that differs between the three *ex-vivo* IGLV6 structures is at position 32, which corresponds to a serine in our structure, a histidine in the 6HUD structure^14^, and an asparagine in the 9OKA structure^21^. In 6HUD, His 32 is in close proximity to Ser 71 and Ser 73, which help coordinate the positively charged histidine side chain within the core of the fibril. In 9OKA, a His residue could be coordinated by the nearby Ser 70 and Asn 72, however its side chain is too large to fit, as suggested by structural clashing determined by ChimeraX, and is thus incompatible with the fold of the 9OKA structure. Instead, Asn 32 forms a favorable hydrogen bonding interaction with Asn 72. In our structure, the hydrophilic Ser 32 is oriented the opposite way and is pointing towards an open, possibly solvated area within the fibril core. Between the three structures, these differences at position 32 are related to the “backbone flipping,” or inversion of the sidechain orientations in these regions which were highlighted previously as a way to preserve the amyloid fold despite differences in amino acids.^21^

Another specific mutation that differs between the three structures is the residue at position 24. Our structure 9ELS and 9OKA present with an arginine at position 24 instead of the glycine found in the 6HUD structure. Glycine substitution in place of the arginine has been shown to decrease stability of the LC native fold, as well as increase the rate of fibril formation.^11,56–58^ In our structure, Arg 24 forms close interactions with many negatively charged and polar residues nearby, namely Asp 85, Asp 88, and Tyr 90. In the 9OKA structure, Arg 24 forms a similar interaction with Tyr 89, and possibly with another ligand within a larger “hydrophilic pore” area.(ref) These interactions stabilize electrostatic forces present as a result of Arg 24 which would otherwise be destabilizing to the fibril. In contrast, Arg 24 appears incompatible with the fibril fold of 6HUD, where a large, charged arginine would impede tight association of the beta strands nearby position 24. In sum, these examples showcase how specific differences in residue composition could contribute to alternative structural conformations.

The available comparison between our structure and the 6HUD and 9OKA structures allows us to speculate a possible folding mechanism for IGLV6-derived fibrils (Fig. 6). All three structures display a ‘C’-shaped turn made up of the double layered peptide backbone spanning approximately residues 13-37 and residues 70-93 (Fig. 6a, blue). This C-shape varies in each fibril structure, with a more angular shape in our structure and a more rounded shape in the 6HUD structure. Additionally, while the N-terminus in our structure is located in a different position from the 6HUD and 9OKA structures (Fig. 6a, red), all three termini have a sharp bend at Pro 14. In our structure, the N-terminus adopts a distal position and wraps around the exterior of the fibril core. In the 6HUD and 9OKA structures, the N-terminus adopts a more proximal position, folding over itself into the interior of the fibril core, previously referred to as a ‘snail shell’-shape.^14,21^ This conformation seems possible in our structure as well, given a large, open channel surrounded by residues 15-40 that could accommodate the N-terminus of our structure.

**Figure 6.**
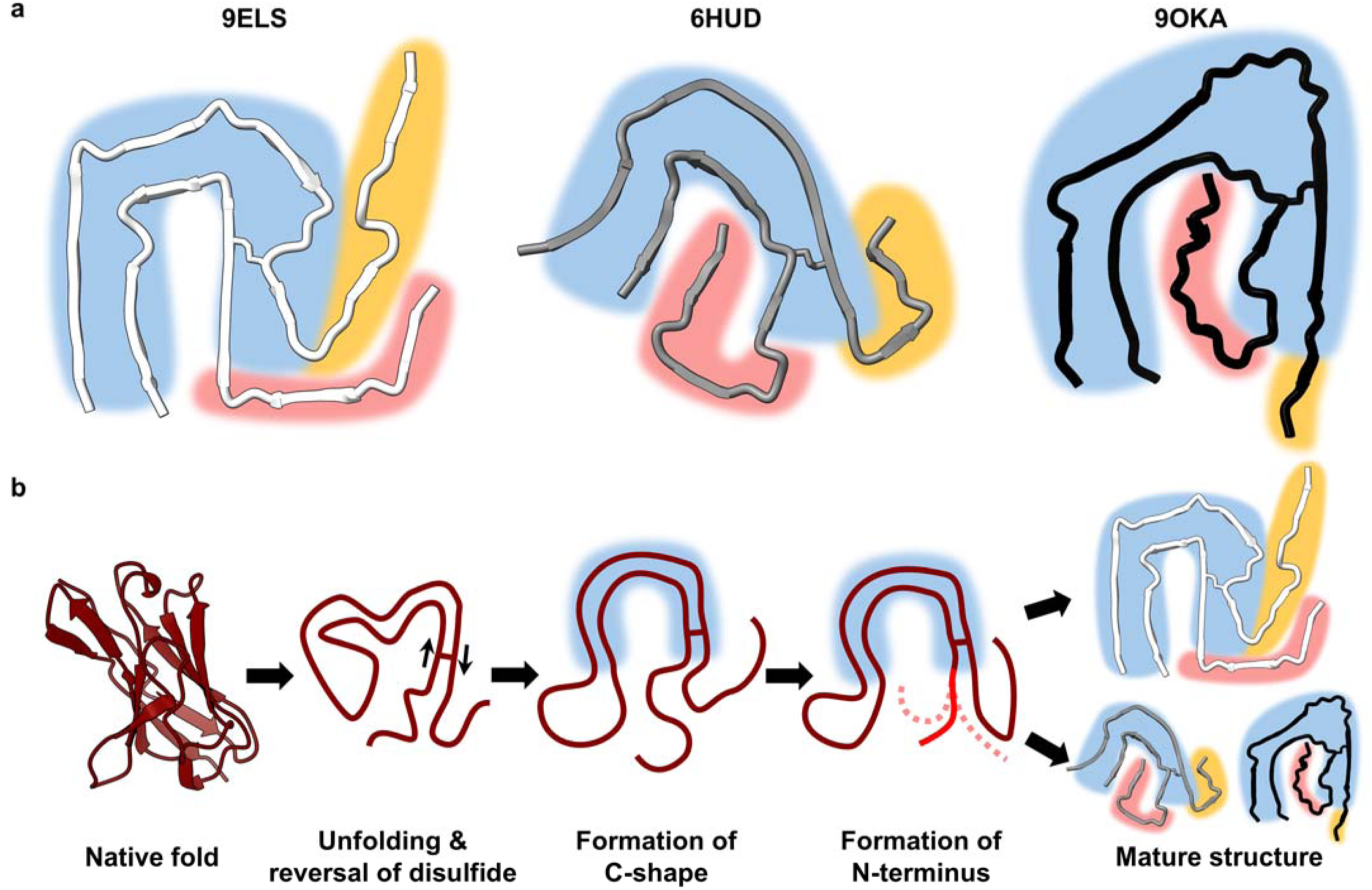
Shared structural features suggest a common folding pathway. **a)** The available IGLV6-derived AL fibrils share certain structural motifs, such as a C-shaped fold (blue) and arrangement of the N-terminal residues (orange) within the interior or exterior of the fibril core. **b) A** proposed folding pathway for IGLV6-derived AL fibrils. In the beginning of the amyloid formation process, the variable domain completely or partially unfolds, followed by reversal of the backbone strands from parallel to antiparallel around the disulfide bond. Then, the common C-shaped motif is formed, followed by divergence of the amyloid folds marked by the differences in N-terminus conformation and further conformational maturation.

Thus, we hypothesize that these common structural motifs between the available IGLV6-derived fibril structures are indicative of a shared folding pathway (Fig. 6b). Following partial or complete unfolding of the variable domain, the strands connected by the disulfide bond would reverse to switch from a parallel to an antiparallel orientation. Then, the C-shaped turn, common to all three structures, would form. Lastly, the fibril structures would diverge from the shared folding pathway as the N-terminus adopts an interior or exterior position in the fibril core. This divergence could be the result of specific residue differences, such as the previously mentioned Tyr 100, or could be the result of interactions with other cofactors found within the tissue environment. Additionally, it is possible to speculate that residues outside the fibril core, such as those found in the constant domain, may have an effect during the folding process. While our data suggests the fibrils consist of truncated LCs without much of the constant domain, some constant domain fragments may linger near fibril deposits or be incompletely removed from the fibrils, consistent with previous studies.^59,60^ In sum, these shared structural motifs could explain part of the mechanism of disease in AL amyloidosis, though more data is needed to support this hypothesis.

Finally, we want to take this opportunity to reflect on cryo-EM, a powerful tool for studying amyloid structures that nonetheless has important limitations worth considering. Bias can be introduced at several stages of the process. Fibril extraction from tissue may preferentially yield species that are more compatible with the buffers and aqueous elution conditions used and may displace cofactors that affect the structures *in vivo*. Cryo-grid optimization tends to favor conditions that produce well-dispersed fibrils, thereby selecting against those that clump or laterally associate. Particle picking in RELION using Topaz is influenced by the hand-picked training set, which may not capture the full ensemble of fibril morphologies. Subsequent class averaging inherently emphasizes the dominant species within the dataset. In our case, two density maps were obtained from ∼40,000 particles each, starting from over 1.4 million particles after Topaz autopicking. It remains unclear why so many particles fail to align into the final density maps—possibly reflecting a broader polymorphism than can be resolved. Nevertheless, cryo-EM remains at the forefront of high-resolution determination of *ex-vivo* amyloid fibril structures.

In conclusion, we have determined the structure of AL fibrils extracted from the explanted heart of a patient with IGLV6-derived AL amyloidosis. We observed the presence of two morphologies, one with a unique and rare rotational symmetry, which raises new questions about the fibril formation process in AL amyloidosis and contributes to the understanding of the amyloid aggregation of LCs *in vivo*.

## METHODS

### Histology of patient tissue

Fresh-frozen human cardiac tissue was thawed overnight at 4°C and then fixed in 20x volume excess of 10% neutral buffered formalin for 48 hours at room temperature with agitation. After fixation, samples were transferred to 70% ethanol and paraffin processed according to established protocols (Sheehan and Hrapchak 1980). Sections were generated from paraffin blocks for hematoxylin and eosin staining (H&E), Congo Red and Thioflavin-S staining. H&E staining was performed on a Sakura DRS-601 x,y,z robot utilizing Leica-Surgipath Selectech reagents according to Sheehan’s textbook methodology. Congo Red slides counterstained with hematoxylin were evaluated for pathologic amyloid aggregation under bright-field following their preparation. Color was imparted to amyloid by methods also described in Sheehan’s text utilizing 0.1% Congo Red in alcoholic-saline following sensitization of the slides with alkaline-alcoholic-saline (10% NaCl, 0.5% NaOH, and 80% EtOH). Thioflavin-S slides were evaluated for foci consistent with plaque amyloid and vascular amyloid, which fluoresce brilliantly when imaged with ultraviolet excitation (400-440nm) and long-pass emission (470 nm+). Thioflavin-S staining was accomplished utilizing methods described by Guntern, et.al. (Guntern et al. 1992). In brief, paraffin sections were brought to water and prepared for alcoholic Thioflavin-S impregnation by sequential oxidation with 0.25% w/v potassium permanganate, bleaching (1% potassium metabisulfite, 1% oxalic acid), peroxidation (1% hydrogen peroxide, 2% sodium hydroxide), and acidification (0.25% acetic acid), all with interceding water washes. Slides were then brought to 50% ethanol and incubated in alcoholic Thioflavin-S (0.004%). After seven minutes staining interval, slides were passed through alcoholic rinses to remove excess Thioflavin-S, dehydrated, cleared, and cover slips affixed with Cytoseal 60 permanent non-fluorescent mounting media (Epredia, Kalamazoo, MI).

### Extraction of AL fibrils

Amyloid fibrils were extracted from flash-frozen tissue as previously described^22^. Briefly, ∼200 mg of frozen tissue was thawed at room temperature and cut into small pieces with a scalpel. The minced tissue was resuspended into 0.4 mL Tris-calcium buffer (20 mM Tris, 138 mM NaCl, 2 mM CaCl_2_, 0.1% NaN_3_, pH 8.0) and centrifuged for 5 min at 5000 × g and 4 °C. The pellet was washed in Tris-calcium buffer three additional times. After the washing, the pellet was resuspended in 0.5 mL of 5 mg mL^-1^ collagenase-Tris-calcium buffer (Type IV *C. histolyticum* collagenase, Sigma-Aldrich) and incubated overnight at 37 °C, shaking at 700 rpm. The resuspension was centrifuged for 30 min at 5000 × g and 4 °C and the pellet was resuspended in 0.4 mL Tris–ethylenediaminetetraacetic acid (EDTA) buffer (20 mM Tris, 140 mM NaCl, 10 mM EDTA, 0.1% NaN_3_, pH 8.0). The suspension was centrifuged for 5 min at 5000 × g and 4 °C, and the washing step with Tris–EDTA was repeated nine additional times. All the supernatants were collected for further analysis. After the washing, the pellet was resuspended in 150 μL of ice-cold water supplemented with 5 mM EDTA and centrifuged for 5 min at 5000 × g and 4 °C. This elution step was repeated three additional times.

### Negative-stained transmission electron microscopy

Amyloid fibril extraction was confirmed by transmission electron microscopy as described.^22^ Briefly, a 2 μL sample was spotted onto a freshly glow-discharged carbon film 300 mesh copper grid (Electron Microscopy Sciences), incubated for 1 min, and gently blotted onto a filter paper to remove the solution. The grid was negatively stained with 2 µL of 2% uranyl acetate for 1 min and gently blotted to remove the solution. Another 5 μL uranyl acetate was applied onto the grid and immediately removed. An FEI Tecnai 12 electron microscope at an accelerating voltage of 120 kV was used to examine the specimens.

### Western Blot

1 µg of fibrils or control was combined with loading dye and heated at 95°C for 10 minutes. The samples were then subjected to electrophoresis on a SurePAGE Bis-Tris 4-12% gel. Recombinant human lambda light chain protein (Abcam) served as the control. The particular recombinant light chain protein used exists in a dimer formation and so appears at twice the normal size of a light chain protein based on characterization of the protein done by the manufacturer. Following electrophoresis, the proteins were transferred onto a 0.2 µm nitrocellulose membrane. The membrane was then probed with a polyclonal rabbit anti-human antibody targeting human AL light chain protein (Abcam, 1:500). A horseradish peroxidase-conjugated goat anti-rabbit IgG (Invitrogen, 1:1000) was used as the secondary antibody. The AL protein was detected using Promega Chemiluminescent Substrate, following the protocol provided by the manufacturer.

### Intact mass spectrometry analysis

20 µL (5 µg) of extracted AL fibrils were spun at 21000 x g for 1hr at 4 °C. The 15 µL of supernatant was removed and 5 µL of 8M GuHCl was added to dissociate the fibrils. Mixture was incubated at 37 °C for 1 h with continuous shaking at 1000 rpm (Eppendorf ThermoMixer). Then, an additional 5 µL of 8M GuHCl was added and incubated at 37 °C for 1 at with continuous shaking at 1000 rpm. Finally, 5 µL of MilliQ water containing reducing agent (DTT) was added. Samples were desalted and analyzed by LC/MS, using a Sciex X500B QTOF mass spectrometer coupled to an Agilent 1290 Infinity II HPLC. Samples were injected onto a POROS R1 reverse-phase column (2.1 x 30 mm, 20 µm particle size, 4000 Å pore size) and desalted. The mobile phase flow rate was 300 μL/min, and the gradient was as follows: 0-3 min: 0% B, 3-4 min: 0-15% B, 4-16 min: 15-55% B, 16-16.1 min: 55-80% B, 16.1-18 min: 80% B. The column was then re-equilibrated at initial conditions prior to the subsequent injection. Buffer A contained 0.1% formic acid in water and buffer B contained 0.1% formic acid in acetonitrile.

The mass spectrometer was controlled by Sciex OS v.3.0 using the following settings: Ion source gas 1 30 psi, ion source gas 2 30 psi, curtain gas 35, CAD gas 7, temperature 300 °C, spray voltage 5500 V, declustering potential 135 V, collision energy 10 V. Data was acquired from 400-2000 Da with a 0.5 s accumulation time and 4 time bins summed. The acquired mass spectra for the proteins of interest were deconvoluted using Bio Tool Kit within Sciex OS in order to obtain molecular weights. Peaks were deconvoluted over the entire mass range of the mass spectra, with an output mass range of 7000-9000 Da, using low input spectrum isotope resolution.

### Protein digestion and enzymatic mass spectrometry

Proteins were precipitated in 23% trichloroacetic acid (Sigma-Aldrich, T0699) at 4 °C overnight. Proteins were pelleted by centrifugation 16100 x g for 30 min at 4 °C. Pelleted proteins were rinsed with 200 μL cold acetone. Air-dried proteins were solubilized in 8 M urea 100 mM Triethylammonium bicarbonate (TEAB) (Sigma-Aldrich, T7408) pH 8.5 and reduced with 5 mM Tris(2-carboxyethyl)phosphine hydrochloride (Sigma-Aldrich, C4706) and alkylated with 55 mM 2-Chloroacetamide (Sigma-Aldrich, 22790). Proteins were digested for both 3 hr and 18 hr at 37 °C in 100 mM TEAB pH 8.5 and varying concentrations of urea (0.5-1.0 M), with 0.5 ug trypsin (Promega, Madison, WI, product V5111) or chymotrypsin (Promega, V106A). Digestion was stopped by addition of formic acid to 5%. (Thermo Scientific). For elastase analysis, 0.5 µg of extracted fibrils were dissolved in a tricine SDS sample buffer, boiled for 2 minutes at 85 °C, and run on a Novex™ 16% tris-tricine gel system using a Tricine SDS running buffer. Gel was stained with Coomassie dye, destained, and the protein smear was cut from the gel. Samples were digested overnight with elastase (Pierce) following reduction and alkylation with DTT and iodoacetamide (Sigma–Aldrich). For proteinase K analysis, 27.25 μg of extracted fibrils were digested for 45 minutes at 37° C while shaking at 400 rpm with 5.45 μg of proteinase K (Thermo Scientific). Digestion was halted with 2 μL of phenylmethylsulfonyl fluoride (PMSF) (Thermo Scientific) before flash freezing. Digested protein was thawed and used for tryptic or chymotryptic digestion and analysis as mentioned above.

Single phase analysis was performed using a Q Exactive HF Orbitrap Mass Spectrometer coupled to an Ultimate 3000 RSLC-Nano liquid chromatography system. Samples were injected onto a 75 µm ID, 15-cm long EasySpray column (Thermo) and eluted with a gradient from 0-28% buffer B over 90 min. Buffer A contained 2% (v/v) ACN and 0.1% formic acid in water, and buffer B contained 80% (v/v) ACN, 10% (v/v) trifluoroethanol, and 0.1% formic acid in water. The mass spectrometer operated in positive ion mode with a source voltage of 2.5 kV and an ion transfer tube temperature of 300 °C. MS scans were acquired at 120,000 resolution in the Orbitrap and up to 20 MS/MS spectra were obtained in the ion trap for each full spectrum acquired using higher-energy collisional dissociation (HCD) for ions with charges 2-8. Dynamic exclusion was set for 20 s after an ion was selected for fragmentation.

### Raw File Processing

The raw data files were converted to .mgf format using MSConvert^61^ GUI. The following filters were applied: 1) “peakPicking vendor msLevel=2-”, 2) “msLevel 2-”, 3) “scanSumming precursorTol=0.01 scanTimeTol=30 ionMobilityTol=5 sumMs1=0”.

Enabling scanSumming is an important feature of this data processing pipeline since the data was acquired with dynamic exclusion disabled. Without dynamic exclusion, the mass spectrometer can repeatedly fragment analyte ions, producing multiple spectra from the same precursor ions. While this ensures that analyte ions are more likely to be fragmented when they are most abundant, resulting in higher quality spectra, it also leads to data redundancy. ScanSumming merges these duplicate spectra, reducing data complexity and improving the signal-to-noise ratio.

### Full MSConvert Command

“msconvert.exe” --mgf --32 --zlib --filter “peakPicking vendor msLevel=2-” --filter “msLevel 2-” --filter “scanSumming precursorTol=0.01 scanTimeTol=30 ionMobilityTol=5 sumMs1=0” --filter “titleMaker <RunId>.<ScanNumber>.<ScanNumber>.<Chargestate> File:“““^<SourcePath^>”””, NativeID:“““^<Id^>”””” “QEX3_1170178.raw” “QEX3_1170180.raw” “QEX2_1181293.raw” “QEX2_1181294.raw” “QEX2_1185163.raw”

### De Novo *Search*

The .mgf files were searched using Casanovo v4.2.0.^25–27^ The trypsin-digested samples were analyzed using the default Casanovo model (*casanovo_massivekb.ckpt*) provided with the v4.0.0 release. Similarly, the chymotrypsin- and elastase-digested samples were analyzed using the non-tryptic model (*casanovo_nontryptic.ckpt*) from the same release. Post-search, each of the result files was individually aligned to ensure a median mass error of 0 ppm for the top 10% of high-scoring peptides. The alignment script used can be found in the GitHub repository.

### Antibody Assembly

Assembly of the antibody sequence from the *de novo* results was done with Stitch v1.5.0.^24^ A customized batch file was created based on the provided monoclonal batch file. All Casanovo input files utilized a CutOffScore of 0.50 and FilterPPM of 5, except for the elastase sample, which used a CutOffScore of 0.85. The FilterPPM and CutOffScore values were determined by analyzing the observed ppm error and score distributions from the top 10% highest scoring peptides for each file. Template matching parameters were configured as follows: EnforceUnique=True, CutOffScore=10, and AmbiguityThreshold=0.9.

During the analysis, Stitch reported a high decoy score, indicating peptides that could be matched to alternative protein sequences. To validate our prediction, the sequence was manually reviewed and later compared to structural data. Three residues in the constant domain (D134, D175, and D176) were picked by Stitch but are uncommon substitutions. These were left as picked by Stitch but may also be asparagines.

An additional joining region template (*>IGLJ2c IGLJ3c*) was added to improve consistency when identifying and filling the gap between the joining and variable regions. This custom template was based on the best matching joining region and several high-scoring peptides which bridged the joining and variable regions. Also, because of the presence of contaminant proteins in the samples, several of the more abundant contaminants identified during the database search were added to the Stitch contaminants file. The custom Stitch files used can be found in the GitHub repository (https://github.com/pgarrett-scripps/LSPB_AL2_Antibody).

### Cryo-EM of AL fibrils

Freshly extracted fibril samples were applied to glow-discharged R1.2/1.3, 300 mesh, Cu grids (Quantifoil), blotted with filter paper to remove excess sample, and plunge-frozen in liquid ethane using a Vitrobot Mark IV (FEI/Thermo Fisher Scientific). Cryo-EM samples were screened on Talos Arctica at the Cryo-Electron Microscopy Facility (CEMF) at The University of Texas Southwestern Medical Center (UTSW). Images were collected using a 300LJkeV Titan Krios microscope (FEI/Thermo Fisher Scientific) with Falcon4i detector operated at slit width of 10 eV at the Stanford-SLAC Cryo-EM Center (S^2^C^2^). Further details are provided in Supplementary Table 1.

### Helical Reconstruction

The raw movie frames were gain-corrected, aligned, motion-corrected and dose-weighted using RELION’s own implemented motion correction program.^62^ Contrast transfer function (CTF) estimation was performed using CTFFIND 4.1.^63^ All steps of helical reconstruction, three-dimensional (3D) refinement, and post-process were carried out using RELION 4.0.^64,65^ The filaments were picked automatically using Topaz in RELION 4.0.^66,67^ Particles were extracted using a box size of 512, 320, and 256 pixels with an inter-box distance of 3 asymmetrical units at helical rise of 4.75 Å. 2D classification of 512-pixel particles was only used to estimate the fibril crossover distance. 2D classifications of 320-pixel particles were used to select suitable particles for further processing. For the single protofilament morphology, a featureless cylinder was used as an initial 3D reference model. For the double protofilament morphology, an initial 3D reference model was generated from a subset of 2D class averages using relion_helix_inimodel2d program, as previously described.^68^ Fibril helix was assumed left-handed for 3D reconstruction. 2D classifications of 256-pixel particles were used for the following refinement steps. 3D auto-refinement was followed by 3D classification without image alignment to filter out suboptimal classes, after which selected particles underwent further 3D auto-refinement. Multiple rounds of CTF refinement, 3D classification (without image alignment), auto-refinement, and post-processing were performed sequentially to reach the highest possible resolution. Final maps were further refined with 3D auto-refinement and were sharpened through post-processing. These maps resulted in a helical rise of 5.03Å, atypical for amyloid fibrils. We believe this inflated value is due to imperfections in the calibration of the electron microscope used for data collection. Overall resolutions were calculated from Fourier shell correlations at 0.143 between two independently refined half-maps using a soft-edged solvent mask.^69^ Local resolutions were estimated using RELION’s Local Resolution tool. Additional processing details are available in Supplementary Table 1.

### Atomic model building and structural refinement

An initial backbone was built using alanine residues in the software COOT v0.9.8.93.^70^ From there, the disulfide bond was positioned according to the structural data and residues were altered according to the predicted sequence. We made real space refinements in both COOT and Phenix v1.20.1 (*phenix.real_space_refine*).^71^ We then used MolProbity to validate the model and identify areas for further refinement. We used the DSSP function on the final PDB model in ChimeraX v1.7.1 to predict the location of secondary structure elements. The final model statistics are listed in Supplementary Table 1.

### Rigid body model fitting

Maps were trimmed to just a few layers using the Map Eraser tool in ChimeraX v1.7.1 for ease of fitting.^72^ Fitting the single protofilament morphology into the double protofilament maps was done using Rigid Body Fit Molecule in COOT v0.9.8.93 for each individual protofilament.^70^

### APR analyses

APRs were predicted using AmylPred2, which is a sequence-based consensus predictor using up to eleven different methods for determining APRs.^73^ At the time of writing, AmylPred2 was unable to access the AmyloidMutants, Aggrescan, and WALTZ webservers. We were able to use Aggrescan^33^ and WALTZ^30^ individually and merge the data with the AmylPred2 results. We were ultimately unable to access the AmyloidMutants webserver and thus used only ten of the eleven methods AmylPred2 usually accesses. We present the data without a consensus threshold to better showcase the breadth of APRs in the 9ELS fibrils. The score for each residue was calculated as the total number of prediction methods that positively predicted a residue as part of an APR. These scores were tabulated for each residue (Supplementary Data 2). The PDB model heat map was generated by replacing the B-factor value in the 9ELS model with the score value from the APR analysis and displayed using ChimeraX v1.7.1.^72^ The sequence heatmap was generated using a custom script.

### Sequence Alignment

To compare the sequence similarity and identity for the 9ELS, 6HUD, 9OKA, and germline LCs, we used the online EMBOSS Needle program via the EMBL-EBI Job Dispatcher server.^41^ The EMBOSS Needle program uses the Needleman-Wunsch algorithm to make pairwise alignments between two sequences.^40^ This gave us individual results for percent identity and percent similarity between the 9ELS sequence and the other LC sequences. For germline comparisons, we used the available protein sequences for the IGLV6-57, IGLJ2, and IGLC2 from IMGT.^74^ For the 6HUD and 9OKA sequences, we used the sequences available via GenBank accession numbers MH670901 and KY432418 respectively. For multiple sequence alignment, we used the CLUSTALW program via the GenomeNET interface, courtesy of the Kyoto University Bioinformatics Center.^75,76^

### Particle tracing

Particles used in helical reconstruction were selected from the output of the 2DClass jobs as class averages. Class averages from RELION corresponding to the desired morphology were selected using subset selection, which resulted in a particles.star file for each particular subset or morphology. We used class averages with a resolution better than 5 angstrom. This cutoff balances a suitable number of particles per micrograph while maintaining a high confidence that the class averages match the intended morphology. Using the CoordinateX and CoordinateY columns within the particles.star file, each individual particle was graphed as a scatterplot and then overlaid onto the corresponding micrograph listed in the MicrographName column.

### Solvation energy calculations

The stabilization energy per residue was calculated by the sum of the products of the area buried for each atom and the corresponding atomic solvation parameters.^29^ The overall energy was calculated by the sum of energies of all residues, and assorted colors were assigned to each residue, instead of each atom, in the solvation energy map.

### Figure Panels

Figures were created with ChimeraX v1.7.1, COOT v0.9.8.93, Stitchv1.5.0, and ESPript3.0 (https://espript.ibcp.fr).^24,70,72,77^

## DATA AVAILABILITY

The mass spectrometry proteomics data have been deposited to MassIVE (ProteomeXchange). MassIVE accession number: *(available to reviewers upon request)*. *De novo* MS data and results can be found here: https://github.com/pgarrett-scripps/LSPB_AL2_Antibody. The PDB and EMDB codes for the single protofilament morphology are 9ELS and EMD-48160. The PDB and EMDB codes for the double protofilament morphology are 9ELT and EMD-48161.

## CONTRIBUTIONS

Conceptualization: L.S., P.T.B.; Data Curation: P.T.B., P.T.G., J.J.M.; Formal Analysis: P.T.B., B.A.N., P.T.G., J.J.M.; Funding acquisition: L.S., J.L.G.; Investigation: P.T.B., V.S., P.T.G., J.J.M., J.E., S.A., M.P., B.M.E., C.L., Y.A., R.P.; Methodology: L.S., J.J.M., P.T.G., P.T.B.; Project Administration: L.S., P.T.B.; Resources: L.S., J.J.M., J.L.G., L.R.R., G.K., S.C.; Software: P.T.B., P.T.G., J.R.Y.; Supervision: L.S.; Validation: P.T.B., B.A.N., P.T.G.; Visualization: P.T.B.; Writing–original draft: P.T.B.; Writing–review and editing: L.S., P.T.B., B.A.N., J.J.M., J.L.G., L.R.R., G.J.M.

## ACKNOWLEDGEMENTS

We are very grateful to the patient who consented to donate their tissue and made this research possible. We would like to thank Michael Sawaya for his valuable feedback and insights on our structures. We would like to thank Douwe Schulte for his assistance in reviewing the *de novo* peptide sequence. We would like to thank Olga Gursky, Chad Hicks, and the other authors of Hicks et al.^21^ for sharing their 9OKA structure with us during our analysis. Additionally, we would like to thank the UTSW Proteomics Core facility for assistance with proteomics experiments, as well as the UTSW Cryo-Electron Microscopy Facility, the UTSW Electron Microscopy Core Facility, and the national cryo-EM facilities Stanford-SLAC (project CA60) for instrumentation, technical support, and data collection.

## CONFLICTS OF INTEREST

L.S. reports current research funding from NHLBI grants DP2HL163810 and R01HL177670, UTSW endowment, UTSW Biotech+ at Pegasus Park Milestone Commercialization Award, and AstraZeneca. L.S. also reports honoraria from Pfizer, and is the founder of AmyGo Solutions. B.A.N. and R.P. are also founders of AmyGo Solutions. J.L.G reports honoraria from Pfizer, NovoNordisk, Tenax Therapeutics, Ultromics, Astra-Zeneca, Alexion, Alnylam, and Eidos/BridgeBio; research funding from Pfizer, Eidos/BridgeBio, Texas Health Resources Clinical Scholars Fund, and NHLBI R01HL160892 and R01HL172993. L.R.R. reports consulting fees from Astra-Zeneca, Eidos/BridgeBio, and Alnylam. The remaining authors declare no competing interests. J.R.Y. reports shareholder and consultant relationships with Chaparral Labs and 3D BioAnalytix, scientific advisory board positions with Yatiri Biotherapeutics, Partnership for Clean Competition, CAS Life Sciences, Mobilion, and OMASS Therapeutics, and Editor in Chief of the Journal of Proteome Research.

## Notes

### Summary of Updates

Information has been expanded and updated. Figures are updated and expanded. Revised manuscript in preparation for submission.

