## Supplementary figures and images for "Polymorphic IGLV6-57 AL amyloid fibrils and features of a shared folding pathway"

### Supplemental Material

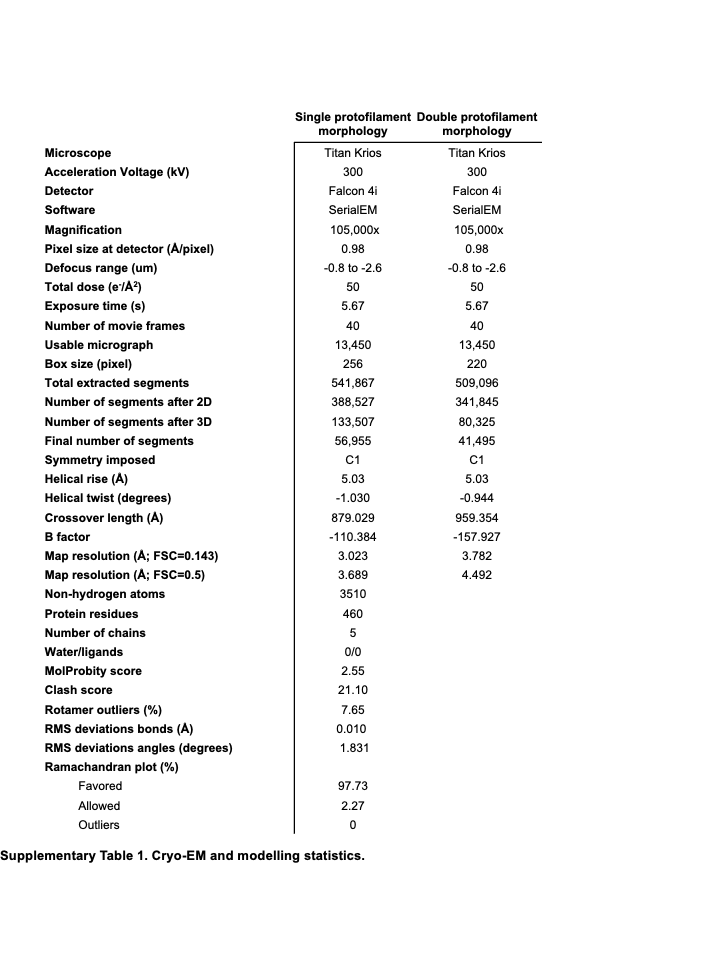

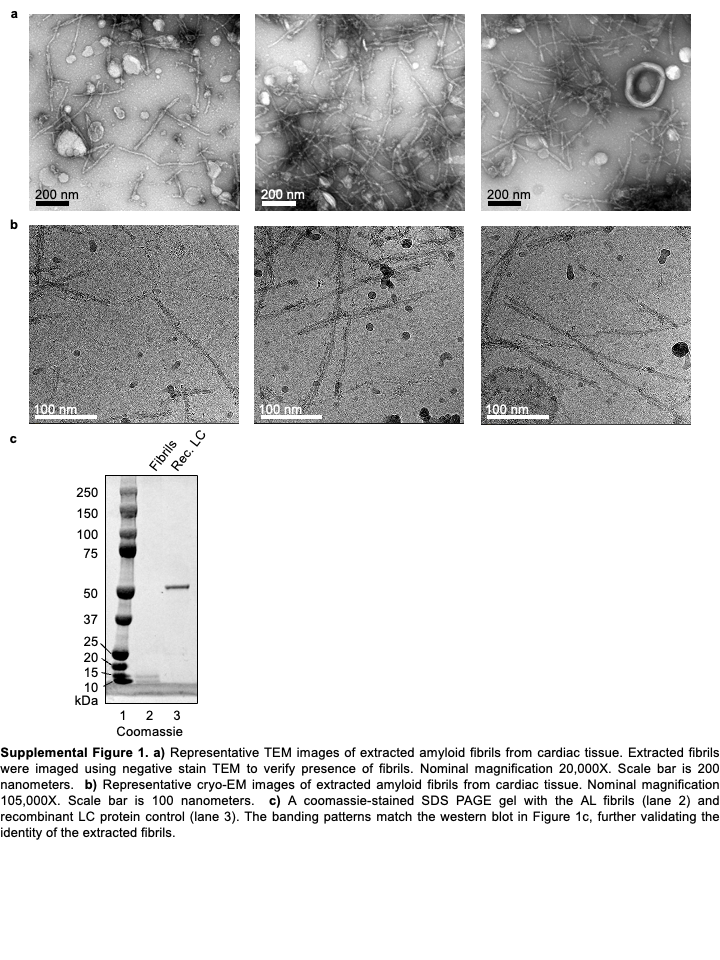

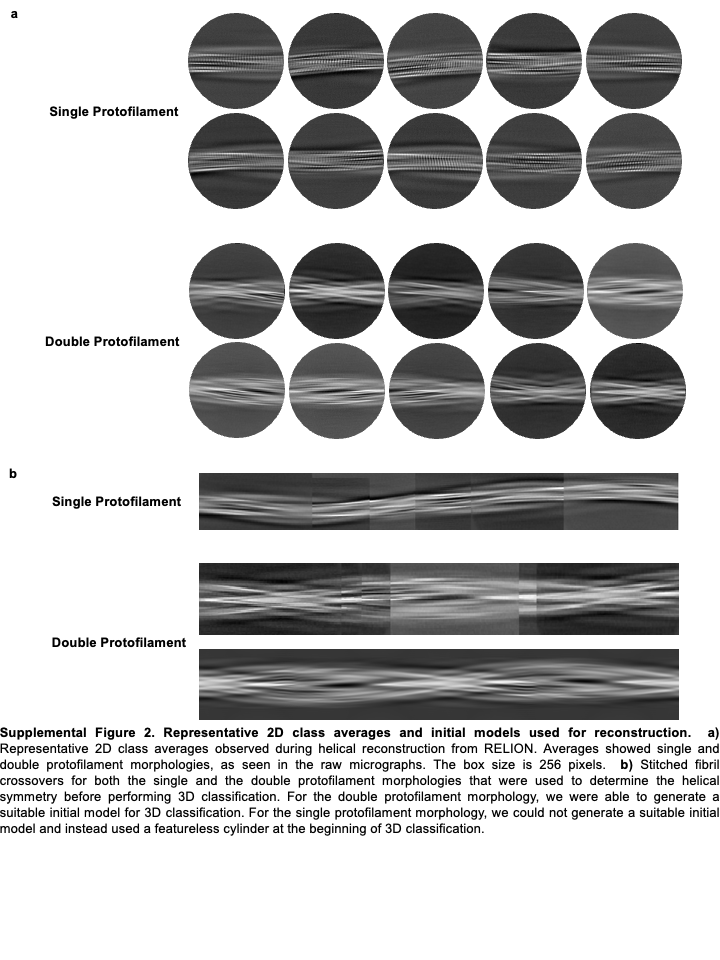

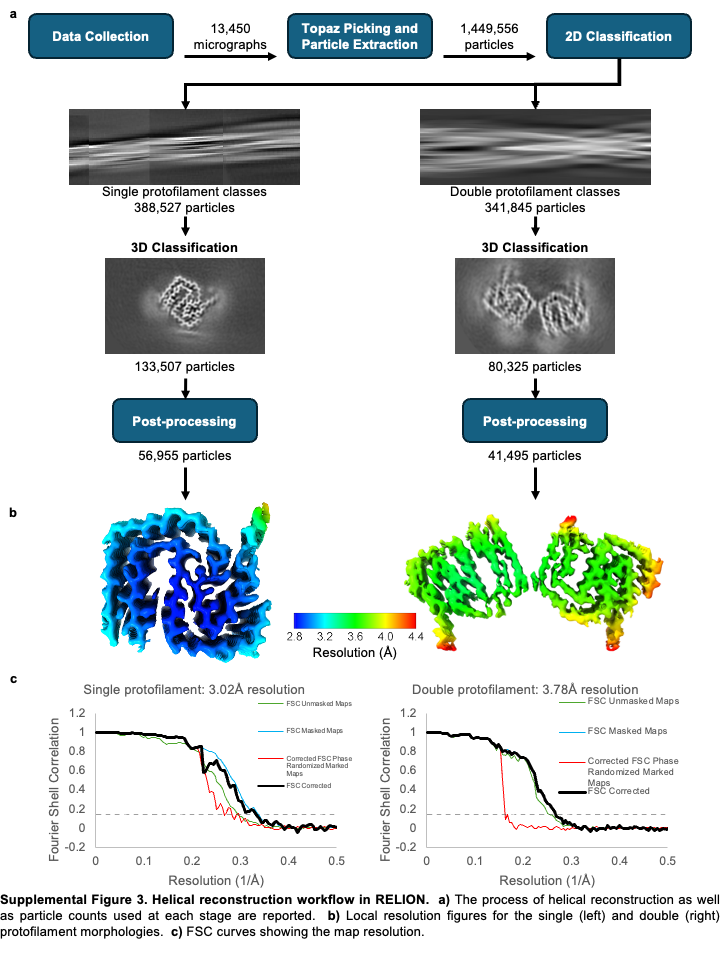

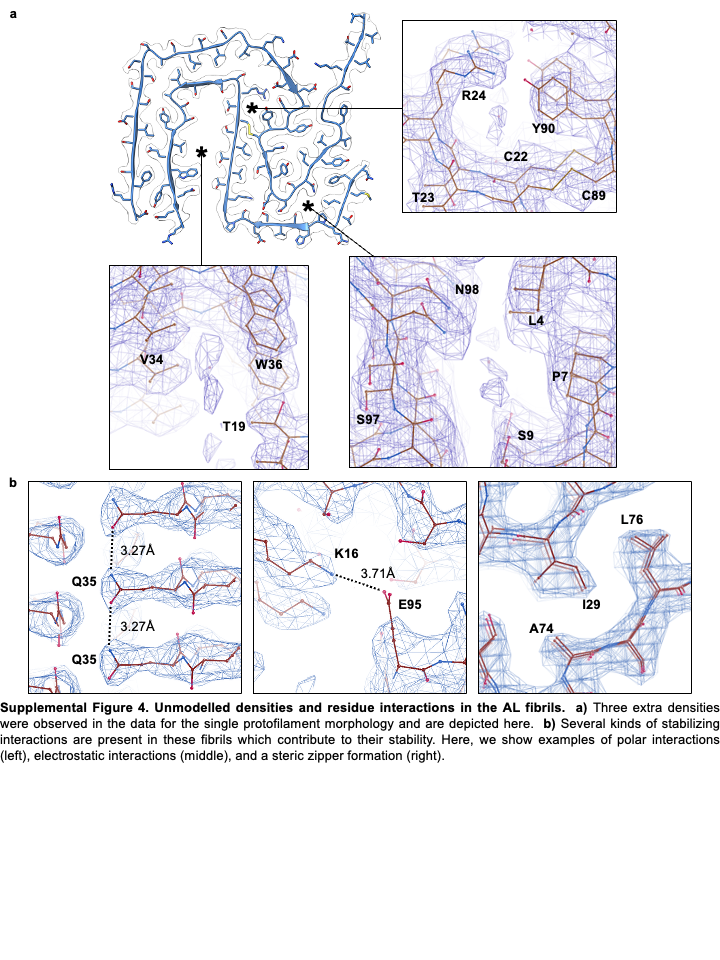

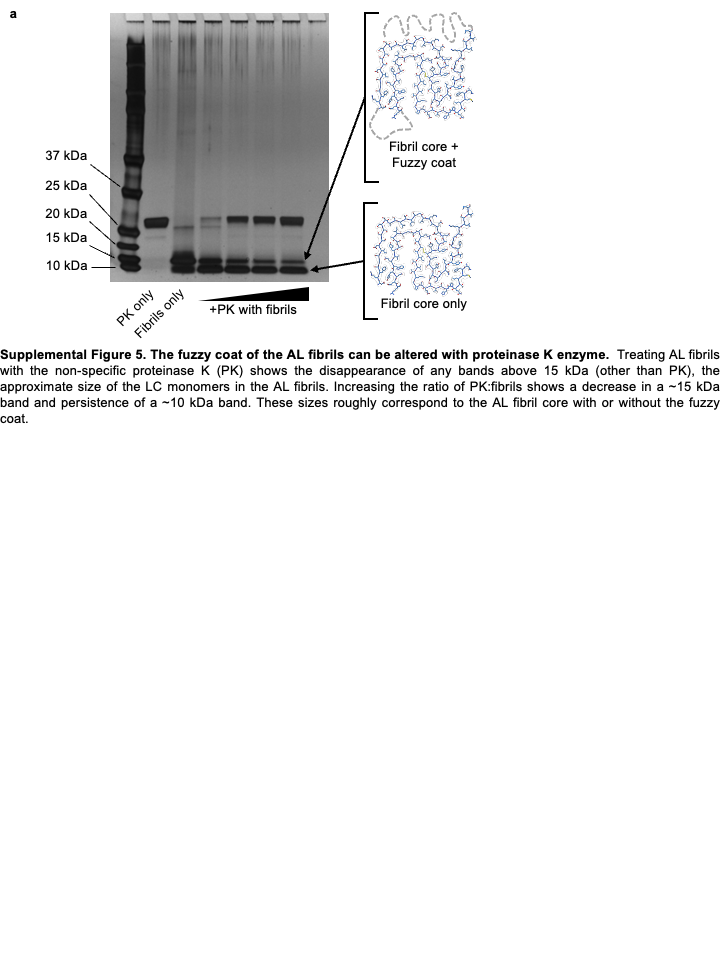

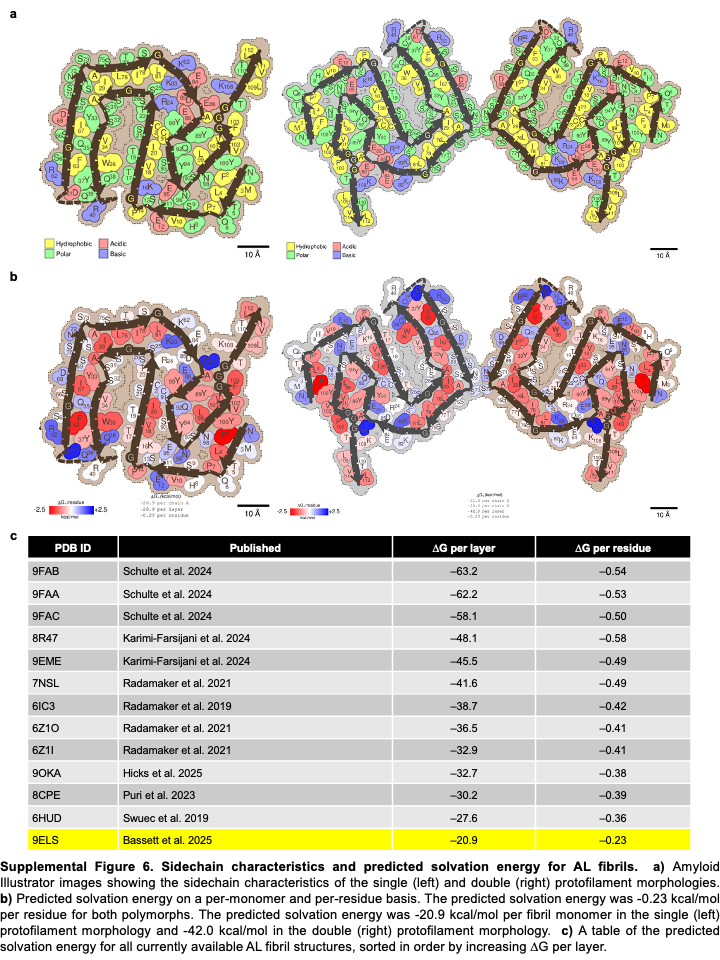

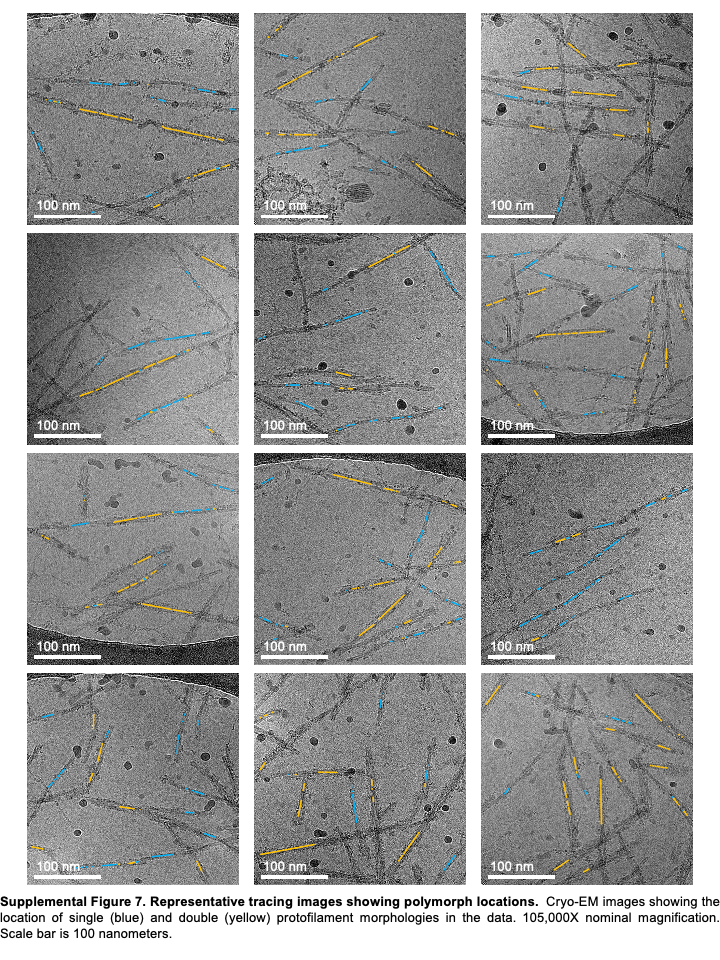

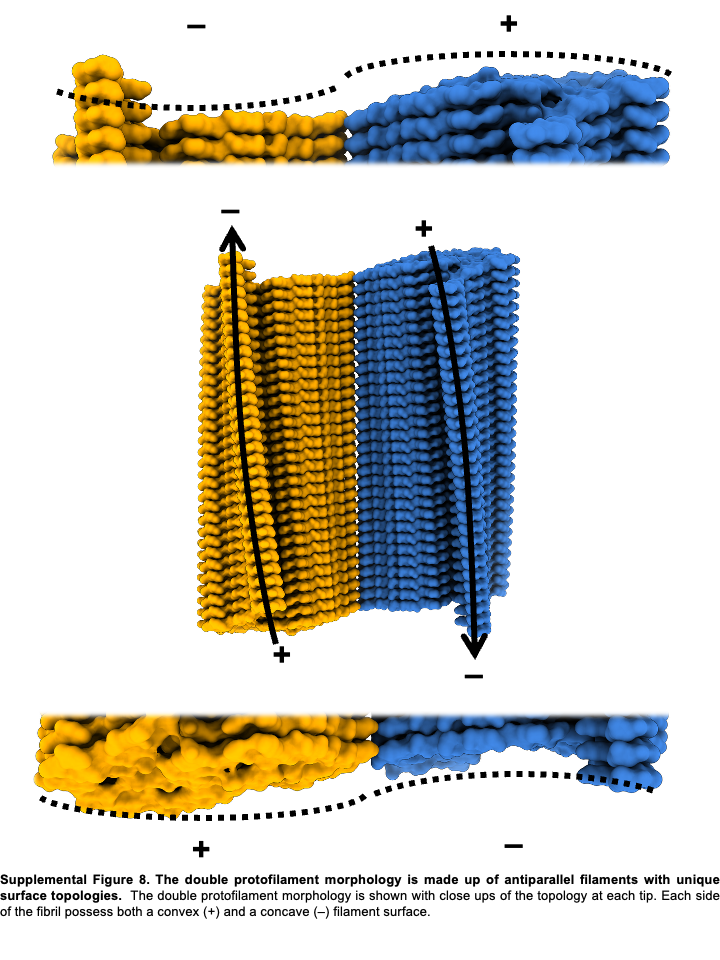
